# The beta-triketone, nitisinone, kills insecticide-resistant mosquitoes through cuticular uptake

**DOI:** 10.1101/2025.01.29.635148

**Authors:** Zachary Thomas Stavrou-Dowd, George Parsons, Clair Rose, Faye Brown, Rosemary Susan Lees, Álvaro Acosta-Serrano, Lee Rafuse Haines

## Abstract

The efficacy of numerous vector control initiatives is compromised by growing insecticide resistance among disease-transmitting arthropods of agricultural, veterinary, and public health significance. Previous investigations on hematophagous (blood-feeding) arthropod vectors, including mosquitoes, have indicated that ingesting blood containing inhibitors of the second enzyme in the tyrosine metabolism pathway, 4-hydroxyphenylpyruvate dioxygenase (HPPD), results in high insect mortality. Building upon this foundation, we evaluated the insecticidal efficacy of the HPPD inhibitor, nitisinone, against susceptible and pyrethroid-resistant strains of three mosquito species: *Anopheles gambiae, Aedes aegypti and Culex quinquefasciatus*. These mosquitoes are vectors of historical diseases such as malaria, emerged diseases such as Dengue and Zika and emerging viral diseases such as the Oropouche and Usutu viruses. We demonstrate, by employing standard screening assays designed to assess the cuticular uptake of mosquitocidal agents, that nitisinone has mosquitocidal activity when blood-fed mosquitoes contact a nitisinone-coated surface. Notably, there is no discernible disparity in susceptibility to nitisinone between an insecticide-susceptible strain of *Anopheles gambiae* and two strains carrying multiple insecticide-resistance mechanisms. We conclude that the mosquitocidal mode of action of nitisinone differs from any of the current 37 classes of insecticides as none have a mode of action that specifically interferes with blood digestion. By highlighting the efficacy of nitisinone as a contact-based insecticide, our findings support the potential expansion of vector control strategies where nitisinone is incorporated into classic interventions like treated bednets and indoor residual spraying.

## Introduction

Exposure to insecticides by the agricultural industry or during public health campaigns for vector control, place intense selective pressure on insect populations and thereby contribute to the emergence of insecticide-resistance and compromises the efficacy of insecticide-based control strategies. Developing new or repurposed chemistries with different insecticidal modes of action is vital to combating this resistance. Pre-2017, malaria cases notably declined, with a global reduction of 18%, but 20% specifically in malaria-endemic countries in Africa. This reduction in malaria incidence is largely attributed to the expansion of key vector control interventions, notably the widespread distribution and use of long-lasting insecticidal nets (LLINs) and indoor residual spraying (IRS) ^1–3^.

However, a reported plateau of malaria cases since 2017 is in part due to mosquitoes’ developing resistance to insecticides. The use of IRS is predicated on the feeding behaviour of the *Anopheles gambiae sensu lato* as this species both feeds and rests indoors. When this mosquito rests on an insecticide-sprayed surface, it dies ^4^. According to the World Health Organization, the most significant effect of IRS occurs after mosquitoes have fed on blood. When resting on an insecticide-treated surface post bloodmeal, a lethal insecticide dose is absorbed through their cuticle, which will immediately compromise the mosquito and prevent malaria transmission ^4^.

*Culex quinquefasciatus*, a global vector of several arboviral diseases, avian malaria and lymphatic filariasis, has also developed resistance to multiple insecticides in countries spanning four continents ^5–13^. Blood feeding is highly dependent on host availability. This species feeds regularly on birds, domestic animals and less often on humans ^14–18^.

*Aedes aegypti* transmits many arboviruses including Dengue, Zika and Chikungunya and is also an urban pest species. This species’ urban nature allows for unconstrained proliferation, particularly as the global population becomes more urbanised. Increasing global temperatures are predicted to further multiply its density and range ^19^. For Dengue virus, these factors have put over half the global population at risk ^20^.

Mosquitoes, as with all hematophagous arthropods, typically ingest a large volume of blood when feeding. This volume can often exceed several times the female’s body weight in a single feeding. For example, mosquitoes, sand flies and reduviid bugs can ingest 3-10 times their body weight in blood ^21^. Mammalian blood is rich in protein and several amino acids, including tyrosine ^22^. One key detoxification enzyme within the tyrosine pathway, 4-hydroxyphenylpyruvate dioxygenase (HPPD), can be blocked using commercially available HPPD-inhibiting herbicides, including members of the b- triketone family ^23^. The repurposed human drug nitisinone ^24,25^, a potent HPPD-inhibitor of herbicides, has been demonstrated to kill several blood feeding arthropods when co-administered with the bloodmeal ^19,26–28^. However, the use of these inhibitors in this manner, either as ectocides or endectocides, could carry ethical concerns and a high safety bar for implementation ^29^.

When mosquitoes take a bloodmeal, their tolerance to pyrethroids naturally increases ^30–34^, which is crucial when selecting IRS products for population control. Both wild Anopheline mosquitoes in various bioassays ^35^ and hut trials ^34^, as well as lab-reared mosquitoes ^31,32,36,37^, have shown reduced mortality after blood feeding compared to non-blood-fed counterparts. Similarly, mosquitoes feeding through baited nets exhibit reduced mortality ^33^. This phenomenon may be linked to the upregulation of enzymes involved in insecticide detoxification ^36^.

Interestingly, standard WHO bioassays only use sugar-fed mosquitoes to validate discriminating concentrations of insecticides ^32^, which may not be lethal to blood-fed mosquitoes. This highlights the importance to consider the effective dose may vary between blood-fed and non-blood-fed mosquitoes, thus impacting residual efficacy and selection for resistance. Additionally, discriminating doses (DD) established for sugar-fed mosquitoes may underestimate resistance levels.

Here we focus on testing three mosquito species, *An. gambiae*, *Ae. aegypti* and *C. quinquefasciatus*, in blood-fed contact assays that mimic landing on walls and targeted for IRS. All blood-fed females are killed when exposed to surfaces coated with nitisinone, but not to other beta-triketone HPPD inhibitors. By leveraging tarsal uptake of HPPD inhibitors, this approach offers a promising strategy for overcoming insecticide resistance and improving vector control. This work justifies further investigation on developing nitisinone for IRS as an alternative to current insecticidal sprays.

## Results

### Mosquitoes must ingest blood to activate HPPD inhibitor-associated killing after cuticular uptake

Sugar-fed, female *An. gambiae* (Kisumu) were tested on tarsal assays. Very low mortality was observed even at the highest dose tested (125 mg/m^2^) for all HPPD-inhibitors screened (Figures 1A and 1B). The corrected mortality at 72 hours post-exposure was nitisinone (5.78%), mesotrione (5.25%), sulcotrione (1.31%) and tembotrione (0%).

**Figure 1.**
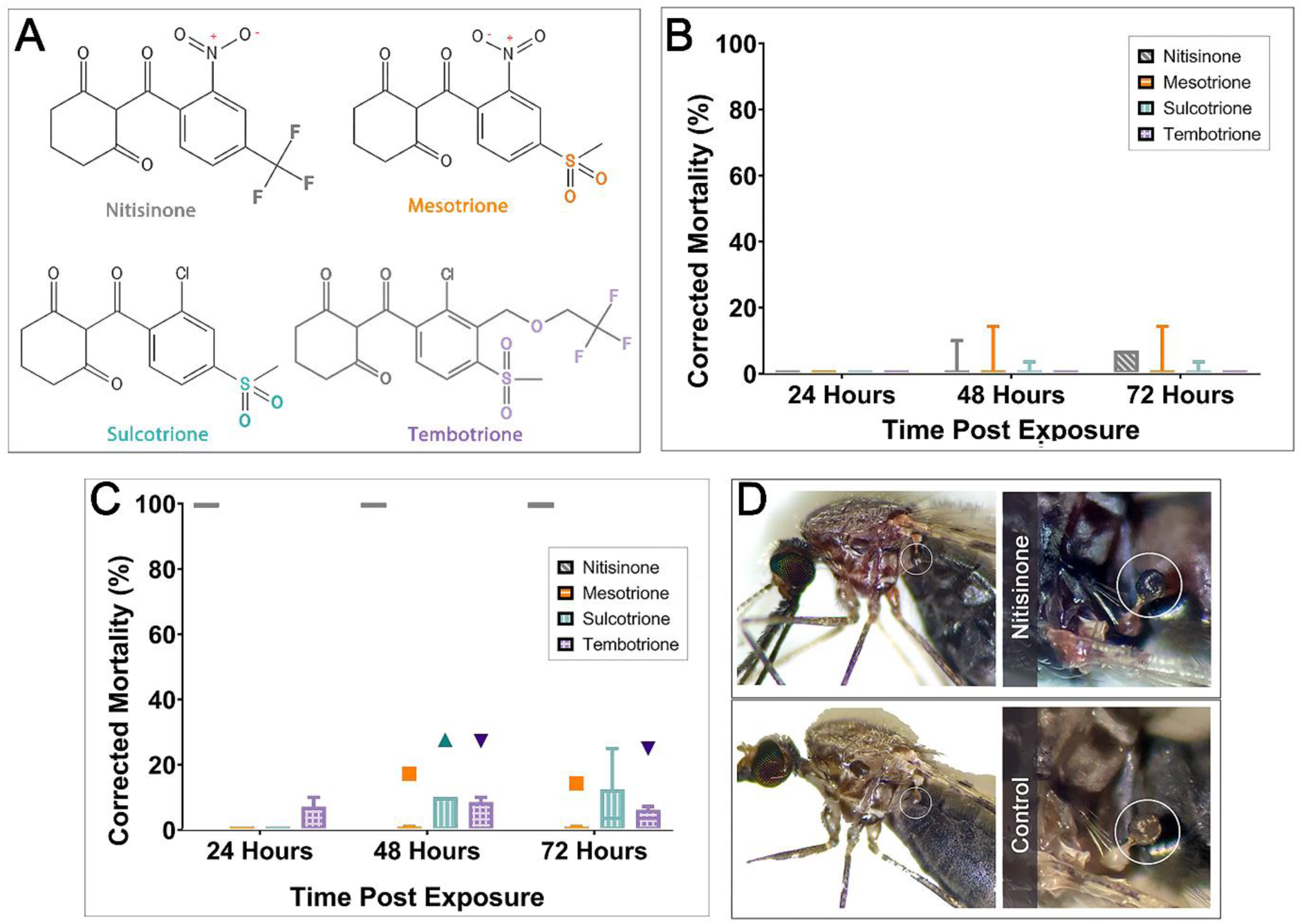
Comparison of mosquito mortality following tarsal contact with a high concentration of HPPD inhibitors (125 mg/m^2^). (A) The four beta-triketones contain a triketone backbone and variable residues on the phenyl group. (B) Sugar-fed *An. gambiae* Kisumu tested on plates coated with the HPPD inhibitors: nitisinone (n = 93) (grey), mesotrione (n = 92) (orange), sulcotrione (n = 98) (turquoise) and tembotrione (n = 95) (purple); all concentrations were at 125mg/m^2^. Time post-exposure is shown as 24, 48 and 72 hours. (C) Blood-fed *An. gambiae* Kisumu survival upon tarsal exposure to HPPD inhibitors: nitisinone (n = 90) (grey)), mesotrione (n = 96) (orange), sulcotrione (n = 93) (turquoise) and tembotrione (n = 96) (purple) 125mg/m^2^. Time post-exposure is shown as 24, 48 and 72 hours. (D) Exposure to nitisinone causes paralysed mosquitoes to systemically darken. The halteres (shown within the white circles) are particularly noticeable.

However, when mosquitoes were **fed blood** prior to (or post bloodmeal (Figure S1)) tarsal exposure, only nitisinone was significantly mosquitocidal (p<0.0001) (Figure 1C) despite the highest inhibitor concentration at 125 mg/m^2^ used for all inhibitors. None of the other inhibitors reached the 80% mortality breakpoint (defined previously as a measure of adequate efficacy of a product ^38^; even after 72 hours post-exposure, sulcotrione (13.24%), tembotrione (11.06%) and mesotrione (5.35%) failed to kill. Of note, exposure to nitisinone caused paralysis in all mosquitoes and they darkened in colour (Figure 1D).

### Topical application of nitisinone on *Anopheles gambiae* shows high intrinsic activity

A single volume of 0.2 µl of each nitisinone dilution was topically applied to the thorax of blood-fed mosquitoes and mortality scored over 48 h. Over 50% killing at concentrations of 0.001 % nitisinone (0.01 mg/ml, representing a dose of 2 ng/mosquito) and above was observed (Figure 2A). Only 0.0001% nitisinone (0.2 ng/mosquito) was not significantly different from the negative control. The breakpoint of 80% was met with the 0.01% (20 ng/mosquito) solution within 24 hours. Collectively, this shows evidence of the high intrinsic activity (inherent potency) of the cuticular exposure of nitisinone in blood-fed mosquitoes.

**Figure 2.**
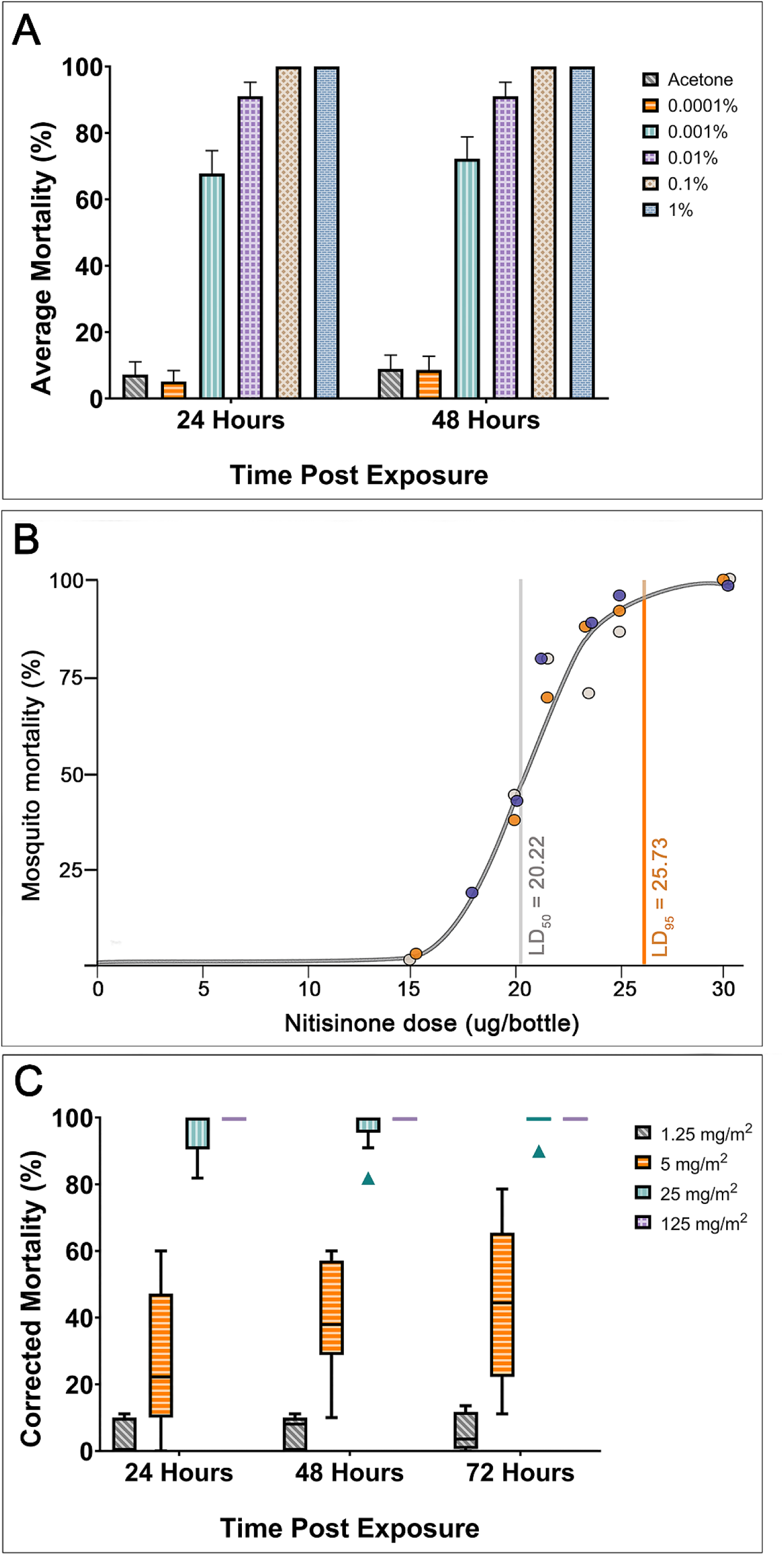
Mortality of female *An. gambiae s.s.* Kisumu to topical, bottle and tarsal assays with nitisinone. (A) Topical assay using blood-fed *An. gambiae* Kisumu tested against a dilution series of nitisinone (three replicates, n =180). (B) Bottle assay testing several concentrations of nitisinone. Rep 1 (grey), Rep 2 (purple), Rep 3 (orange). LD_50_ (grey vertical line) is the dose of nitisinone killing 50%. LD_95_ (orange vertical line) is 95% of the population is killed at this dose. (C) Tarsal assay on blood-fed mosquitoes exposed to four different nitisinone concentrations (1.25 – 125 mg/m^2^).

### Discriminating dose of Nitisinone

As nitisinone was the only mosquitocidal HPPD inhibitor in the tarsal and topical bioassays, we continued to follow the testing cascade ^38^ using the WHO Bottle Assay to establish a discriminating dose as a measure of potency (Figure 2B). Blood-fed *Anopheles gambiae* Kisumu were exposed to nitisinone-coated bottles in a modified CDC Bottle Assay ^38^ and the dose-response graph is shown (Figure 2B). The LD**_50_** of nitisinone was calculated at 20.22 µg/bottle (95% CI 19.55 – 20.69), which is equivalent to 0.72 mg/m^2^. The LD**_95_** was calculated at 25.73 µg/bottle (95% CI 24.65 – 27.37), which is equivalent to 0.92 mg/m^2^. The discriminating dose is calculated as three times the LD**_95_** and is therefore 77.19 µg/bottle, which is equivalent to 2.76 mg/m^2^.

### Insecticide-resistant strains of *An. gambiae* are killed by tarsal uptake of nitisinone

Different strains of insecticide-resistant and susceptible *Anopheline* mosquitoes were tested for susceptibility to nitisinone upon tarsal contact. The resistant mosquito strains were selected because of their well-documented resistance mechanisms to several insecticide classes and carefully documented GLP rearing conditions including maintaining under insecticide selection pressure and genotyping for contamination (Table S1). *An. gambiae* Kisumu (susceptible, Figure 2C), Tiassalé 13 (resistant, Figure 3A) and *An. coluzzii* VK7 2014 (resistant, Figure 3B) all died within 24h of exposure to nitisinone at 125 mg/m^2^. At a 5-fold lower dose, 25 mg/m^2^, the killing effect was slightly reduced (insignificant) although it remained above the 80% threshold by 72h post exposure: Kisumu (98.9 %), Tiassalé 13 (92.5%) and VK7 2014 (92.9%). At lower nitisinone doses (5 mg/m^2^ and 1.25 mg/m^2^) the mortality for all timepoints was below the 80% threshold, suggesting the cut-off for this assay was 25 mg/m^2^ for all strains tested. At 72 hours there was no significant difference between the strains at any concentration except at 1.25 mg/m^2^ between Kisumu (which was significantly lower (p = 0.003) than VK7 2014). Comparing how long it took for the mosquitoes to die post-exposure, there is no clear pattern (Figure 3C). However, at the lowest dose, Kisumu reaches its maximum mortality quicker than Tiassalé 13, which is slower to do so at 25 mg/m^2^. It seems that VK7 2014 is slower to reach maximum killing at 5 mg/m^2^ and 25 mg/m^2^. Insecticide-resistant strains of *C. quinquefasciatus* and insecticide susceptible strains of *Ae. aegypti* are killed by the tarsal uptake of nitisinone.

**Figure 3.**
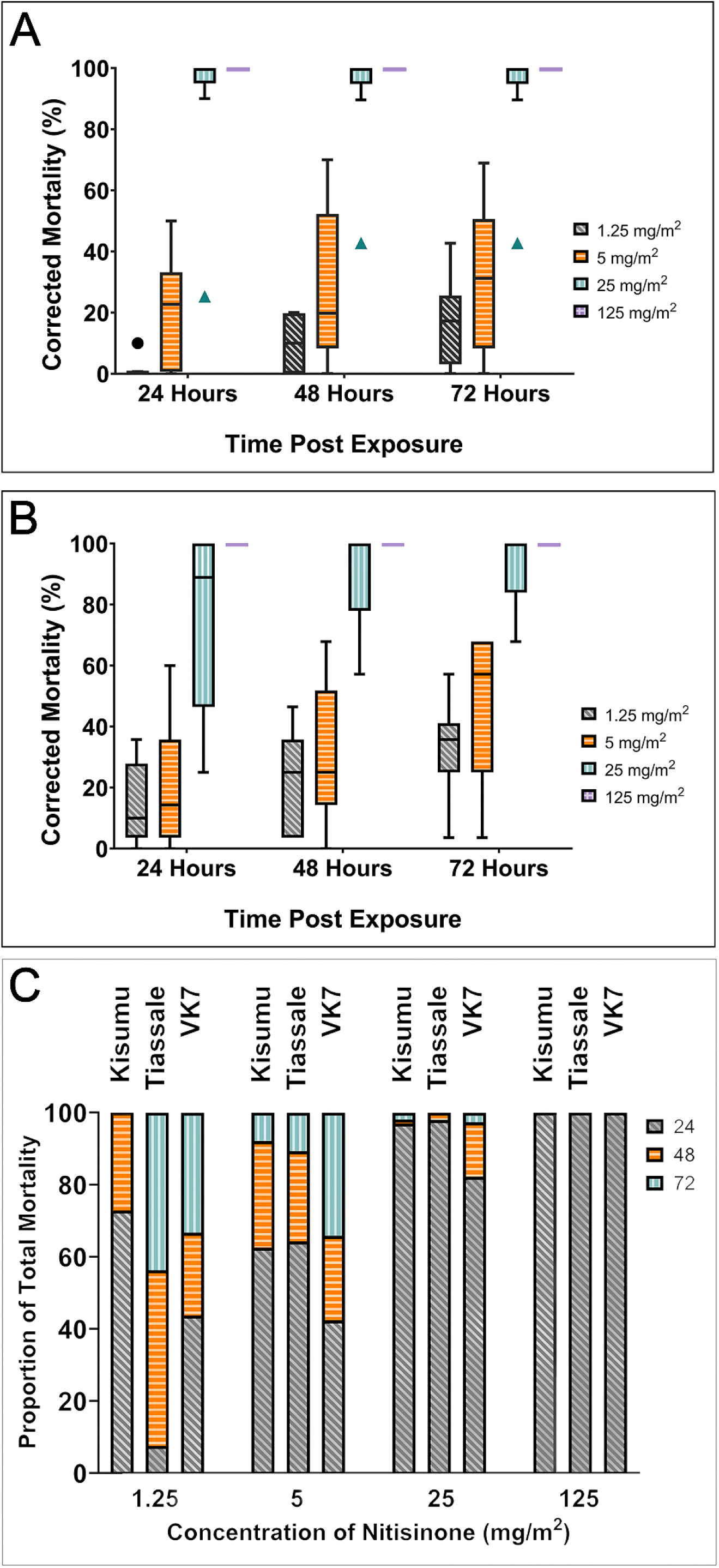
Tarsal assays testing nitisinone efficacy against insecticide resistant *An. gambiae s.l.* Tiassalé 13 (A) and VK7 2014 (B). Time taken for mortality to occur in each species of *Anopheles gambiae* (0 -72 hours) (C) Speed of killing as a proportion of total mortality for each strain of *Anopheles gambiae* tested at each concentration.

To confirm that the tarsal exposure of nitisinone had broad activity against other mosquito species, the insecticide-restraint strain of *C. quinquefasciatus* Muheza (Figure 4A) and the insecticide susceptible *Ae. aegypti* New Orleans (Figure 5B) were exposed to tarsal plates coated with nitisinone. For Muheza at 125 mg/m², there was a breakpoint killing of 81.6% at 24 hours, which increased to 94.7% by 72 hours. This killing effect was slightly lower, though not significant, compared to the 100% mortality observed in the *Anopheles* strains (Figure 5). At 25 mg/m², the killing effect was significantly reduced (<0.0001) to 35.8% by 72 hours. Mortality decreased further at 15 mg/m² (7.9% at 72 hours), 5 mg/m² (1.6% at 72 hours), and 1.25 mg/m² (1.8% at 72 hours), indicating a potentially higher innate resistance to lower concentrations of nitisinone. The reduction in killing efficacy by 72 hours, compared to the Anopheles strains, was significant (p-values ranging from 0.03 to <0.0001) across all comparable doses: 25 mg/m² and 5 mg/m² (15 mg/m² was not tested in Anopheles), and at 1.25 mg/m² for VK7 2014 and Tiassalé 13 (though not significant for Kisumu).

**Figure 4.**
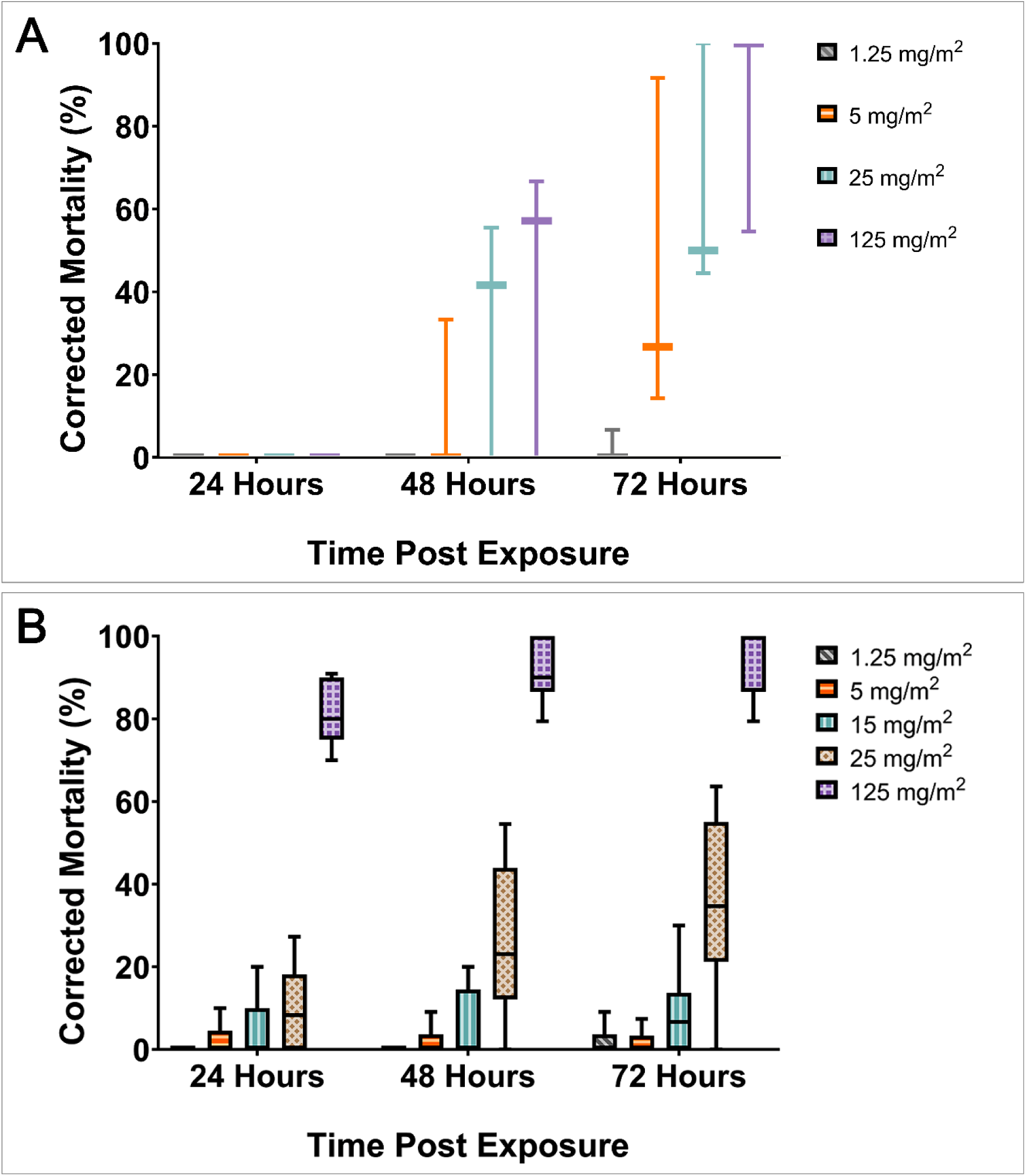
*Culex quinquefasciatus* (A) and *Aedes aegypti* (B) susceptibility profiles to tarsal contact with nitisinone. A) Nitisinone concentrations tested: 1.25 mg/m^2^ (grey), 5 mg/m^2^ (orange), 15 mg/m^2^ (turquoise), 25 mg/m^2^ (brown) and 125 mg/m^2^ (purple). B) 1.25 mg/m^2^ (grey), 5 mg/m^2^ (orange), 25 mg/m^2^ (turquoise) and 125 mg/m^2^ (purple). Three biological replicates represent n = 90 per dose.

**Figure 5.**
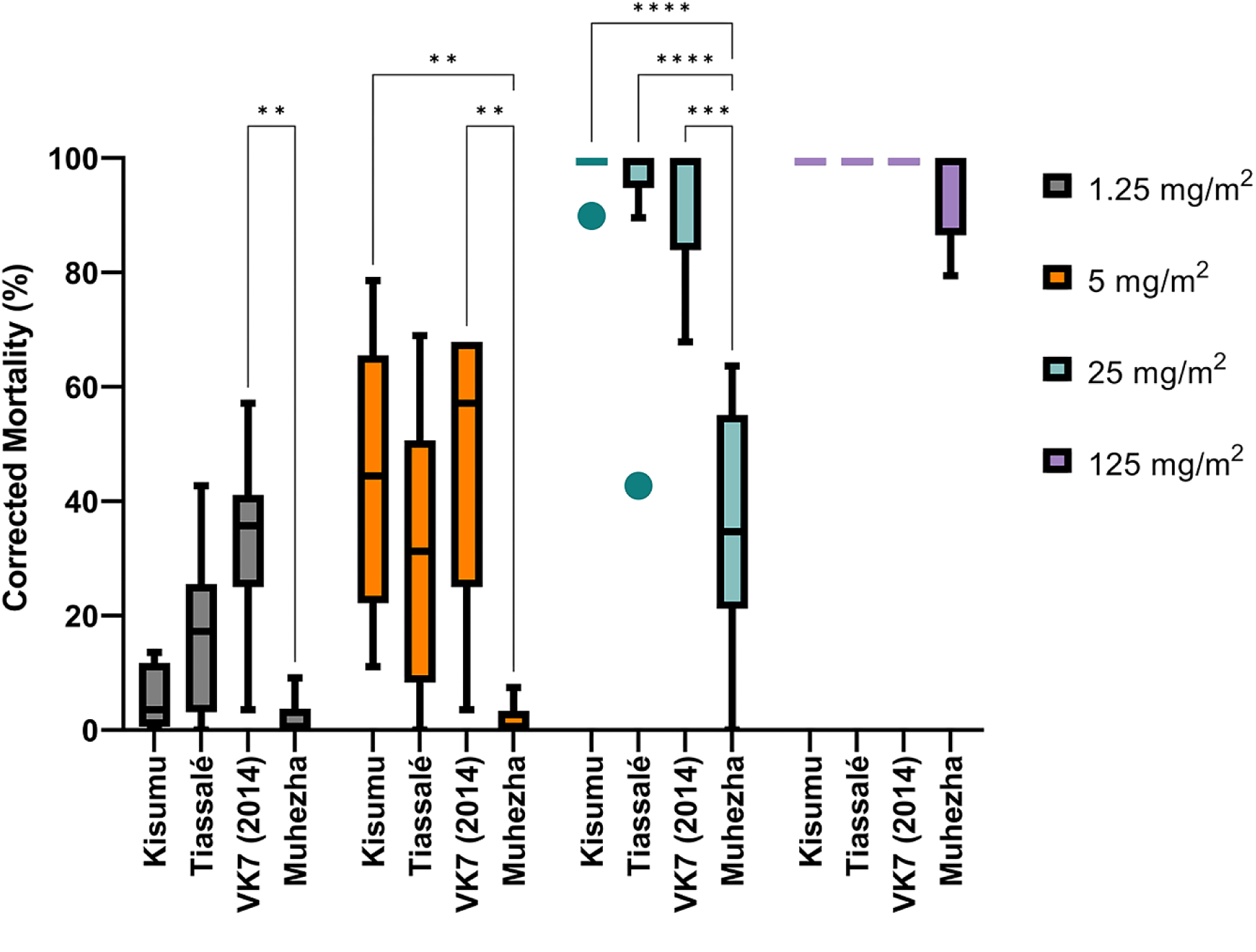
The comparative mortality of four mosquito strains 72 h post-tarsal exposure to nitisinone. Kisumu (insecticide-susceptible *An. gambiae*), Tiassalé 13 (insecticide-resistant *An. gambiae*), VK7 2014 (insecticide-resistant *An. coluzzii*), Muheza (insecticide-resistant *C. quinquefasciatus*) (x-axis). Three biological replicates for each group with an average of 90 mosquitoes were used per replicate (or used for each assay). Nitisinone concentrations were: 1.25 mg/m^2^ (grey), 5 mg/m^2^ (orange), 25 mg/m^2^ (turquoise) and 125 mg/m^2^ (purple).

For *Aedes aegypti* New Orleans (Figure 4B), the potency of nitisinone was reduced compared to both *Anopheles* and *Culex*. The breakpoint killing at 125 mg/m² was not reached until 48 hours (84.8%), rising to 97.0% by 72 hours. No other concentration passed the breakpoint by 72 hours: 25 mg/m² (77.8%), 5 mg/m² (60.3%), and 1.25 mg/m² (6.7%).

However, despite mortality differences at the highest dose between New Orleans and Muheza, by 24 hours, all other concentrations were more potent in New Orleans (susceptible) than Muheza (resistant).

## Discussion

In the pursuit of innovative vector control strategies, a promising avenue to identify new insecticidal compounds is to extend the search beyond conventional neurological and detoxification gene targets to include insect blood-feeding mechanisms. Previously, nitisinone was shown to be either toxic when ingested by blood-feeding insects or when absorbed through the cuticle via topical application (solvent assisted) ^26–28,39^. Adding to previous results that nitisinone induced mortality in tsetse and triatomines, we provide additional evidence that inhibiting 4-hydroxyphenylpyruvate dioxygenase (HPPD) constitutes a viable approach for mosquito control via cuticular uptake. The four beta-triketone HPPD inhibitors we investigated were originally selected on commercial availability and potency on plant targets or, in the case of nitisinone, FDA approval as a human drug ^23,25^. We have shown nitisinone is the only HPPD inhibitor with contact-based mosquitocidal effect. The lack of significant toxicity of mesotrione, sulcotrione and tembotrione was unexpected, but it might reflect differences in the cuticular penetrability among these chemistries. Furthermore, nitisinone killed mosquito strains resistant to several insecticide classes, providing evidence of its potential to control insecticide-resistant mosquitoes. Using the glass plate tarsal assay, there was no difference in mortality at 5, 25 and 125 mg/m^2^ nitisinone between the insecticide-susceptible Kisumu strain and the insecticide-resistant Tiassalé 13 and VK7 2014 strains. This is not surprising as tyrosine metabolism is essential to hematophagous insects and therefore its inhibition should be lethal to blood-feeding arthropods ^39^. We predicted that VK7 2014, and to some extent Tiassalé 13, would show less mortality than Kisumu because of cuticular changes to the tarsi that slow uptake of insecticides (*i.e.* thickness of cuticle) ^40,41^. However, this was not observed, thus indicating tarsi remain equally permeable to nitisinone. It remains to be determined if the sensory appendage protein 2 (SAP2), which has been shown to be compound-specific in resistant mosquitoes, potentially mediates nitisinone uptake across the cuticle rather than a generic cuticular thickening ^42^.

Incorporating data from several types of assays can enhance the robustness of insecticide efficacy evaluations. However, it is essential to acknowledge that the topical application assay is the least representative of field conditions among the three methods discussed. The direct application of insecticide onto the mosquito thorax using an aqueous solution does not mimic a typical environmental exposure scenario ^38^, although it can give a rough indication of arthropod susceptibility to a compound ^28^. Although the glass plate and bottle assays both measure bioefficacy via tarsal contact, their results are not directly comparable. Differences in exposure duration and surface coating significantly influence the mortality rates observed in each assay, thus highlighting the importance of assay selection to accurately assess insecticide efficacy.

Insecticide residual spraying (IRS) exploits the mosquito behaviour to rest post blood-feeding, and consequently adsorb insecticide upon contact with an insecticide-covered surface. The efficacy of IRS can be significantly reduced by insecticide degradation, sub-optimal coverage and modifications to sprayed surfaces such as wall washing post-treatment. These issues cause two problems: 1) mosquito survival post-exposure to a non-lethal dose and 2) exposure to a sublethal dose can drive insecticide resistance ^43^. As we used blood-fed mosquitoes rather than the industry standard of sugar-fed, direct comparisons to previous published data is not possible. However, comparing discriminating dose (DD) and dose response curve shape for nitisinone with data generated for other compounds ^38^ is encouraging. The discriminating dose combines a fixed exposure time, and the amount of insecticide coated inside a glass bottle with the uptake depending on the time of actual tarsal contacts. Based on these results, the potency of nitisinone is greater than clothianidin, spinetoram, metaflumizone and dinotefuran ^38^, which makes it attractive for further optimisation as novel IRS formulations. Considering the gradient of the dose-response curve (roughly determined by calculating the gradient from the LC_95_ and LC_50_ derived from Figure 3), nitisinone has the steepest curve, indicative of high potency. This reflects previous nitisinone results in both blood-feeding and topical application assays in another dipteran disease vector, *Glossina m. morsitans* ^26^. We briefly tested the efficacy of nitisinone (using the glass plate assay) by exposing Kisumu or New Orleans before blood feeding (Figure S1). Nitisinone is still active tarsally and this models a mosquito landing on a sprayed wall before taking a blood meal which warrants further investigation. It is possible that nitisinone’s efficacy (and perhaps that of other HPPD inhibitors) through tarsal contact could be increased by combining adjuvants such as rapeseed oil methyl esters as described with other insecticides ^44,45^. This was carried out using the exposure pre blood feed with Kisumu and showed a significant increase in mortality at the 5 mg/m^2^ concentration (Figure S2).

There are compelling reasons to formulate nitisinone into an IRS: 1) nitisinone targets blood-fed (naturally more insecticide tolerant) mosquitoes resting on sprayed surfaces indoors, suggesting a novel mechanism of action that could potentially overcome increased insecticide-resistance ^4^; 2) it is stable under a range of environmental conditions such as UV radiation, temperature and pH ^26,46^; and 3) it is unlikely to be immediately metabolised by P450 enzymes, although resistance through P450 metabolism could slowly evolve with selective pressure ^26^.

The killing kinetics of unformulated nitisinone across the different insecticide-resistant mosquito strains is intriguing. Strain VK7 2014 may die slower because of cuticular thickening, reduced blood meal size or speed of bloodmeal digestion, which we did not examine. The lower toxicity of nitisinone with the insecticide-resistant *Culex* strain, Muheza, indicates that a higher concentration (between 25 and 125 mg/m^2^) needs further investigation. Additionally, as with *Culex, Aedes* showed reduced susceptibility to nitisinone compared to *Anopheles,* which may indicate physiological differences between the two species in blood-feeding and digestion rates ^27^.

In conclusion, our results show nitisinone kills blood-fed mosquitoes via tarsal contact and mesotrione, sulcotrione, and tembotrione do not. This killing did not discriminate between mosquito strains susceptible or highly resistant to other insecticide classes (including pyrethroids, organochlorides, and potentially carbamates). Moreover, the mosquitocidal efficacy of nitisinone via cuticular absorption extends beyond Anopheline species, as demonstrated by its effectiveness against *C. quinquefasciatus* and *Aedes aegypti*. Our data supports further investigation into optimizing nitisinone uptake, potentially through the use of chemistries to enhance cuticular absorption or by formulating nitisinone with adjuvants. With its novel mechanism of action, nitisinone has the ability to exploit the blood-feeding behaviours of female mosquitoes. This makes it a promising candidate for innovative indoor residual sprays and long-lasting insecticidal nets, particularly in regions where traditional mosquito control methods are being undermined by the rapid emergence of pyrethroid resistance.

## Methods

### Mosquito rearing for bioassays

Three strains of Anopheline mosquitoes aged 3 – 5 days were used for the initial bioassays: *An. gambiae* s.s. Kisumu strain (insecticide susceptible), *An. gambiae* s.l. Tiassalé 13 strain (insecticide-resistant) and *An. coluzzii* VK7 2014 strain (insecticide-resistant). The Kisumu strain was reared in-house at the Liverpool School of Tropical Medicine (LSTM) whilst adult mosquitoes of the Tiassalé 13 and VK7 2014 strains were obtained from the GLP-accredited Liverpool Insect Testing Establishment (LITE) ^45^. In addition, LITE provided a pyrethroid resistant strain of *C. quinquefasciatus* (Muheza) and an insecticide susceptible strain of *Ae. aegypti* (New Orleans) ^47^.

The Tiassalé strain, originating from Côte d’Ivoire, has developed resistance to pyrethroids, DDT and carbamates (Table S1). The VK7 (2014) strain, originating from Burkina Faso, has high resistance intensity to pyrethroids and DDT (Table S1). Both Tiassalé 13 and VK7 2014 are maintained at LITE under deltamethrin selection pressure. Muheza is resistant to the pyrethroids permethrin and deltamethrin (Table S1). Full resistance validation can be gained from LITE ^47^.

The Kisumu strain was reared following previously published methods: 26°C ± 2°C and 80% relative humidity ± 10%, with a 12hr:12hr light: dark cycle that includes an hour of dawn and dusk. Larvae were maintained on fish food (TetraMin® tropical flakes, Blacksburg, VA). To encourage female mosquitoes to deposit eggs, mated females were fed human blood purchased from the UK’s National Blood Authority (https://www.blood.co.uk) using a Hemotek Membrane Feeding System (Hemotek Ltd., Blackburn, UK). Adult mosquitoes were maintained on 10% sucrose solution *ad libitum*.

### Mosquito blood feeding for insecticide-based bioassays

Female mosquitoes were sugar-starved for five hours and then offered human blood using the Hemotek Membrane Feeding System (Hemotek Ltd., Blackburn, UK) *ad libitum* for 45 minutes. After this time, the blood was removed, and fully engorged females were gently aspirated for bioassay screening. In assays using non-blood-fed mosquitoes, the same sugar starvation process was followed.

### Screening of HPPD Inhibitors using the Glass Plate Tarsal Bioassay

Four HPPD inhibitors were chosen for tarsal screening based on previous potency as an ectocide and the availability for purchase as either commercial herbicides or human drugs (Table S2) ^23,26^. All four selected inhibitors were purchased from Merck Life Science, Gillingham, UK.

Using the glass plate bioassay (Figure 6) previously described ^38^, a total of three replicates (biological) of thirty Kisumu mosquitoes (ten female mosquitoes per plate) were exposed to inhibitor-coated glass Petri dishes (VWR, Lutterworth, UK). Briefly, glass plates were coated with a 500 µl solution of either nitisinone, mesotrione, sulcotrione or tembotrione dissolved in acetone (Fisher Scientific, Loughborough, UK) to a concentration of 125 mg/m^2^ (chosen as the highest dose in previous screening work ^38^). The solvent, acetone, was used as the negative control. Petri dishes were dried for 4 hrs at room temperature in a fume hood prior to use. Mosquitoes were exposed to coated plates for 30 minutes and then carefully removed from the testing plate to a cage. Mortality was tracked at 30 min, 24h, 48h and 72h post exposure. The assay was repeated using non-blood- fed mosquitoes (as described above) as the negative control.

**Figure 6.**
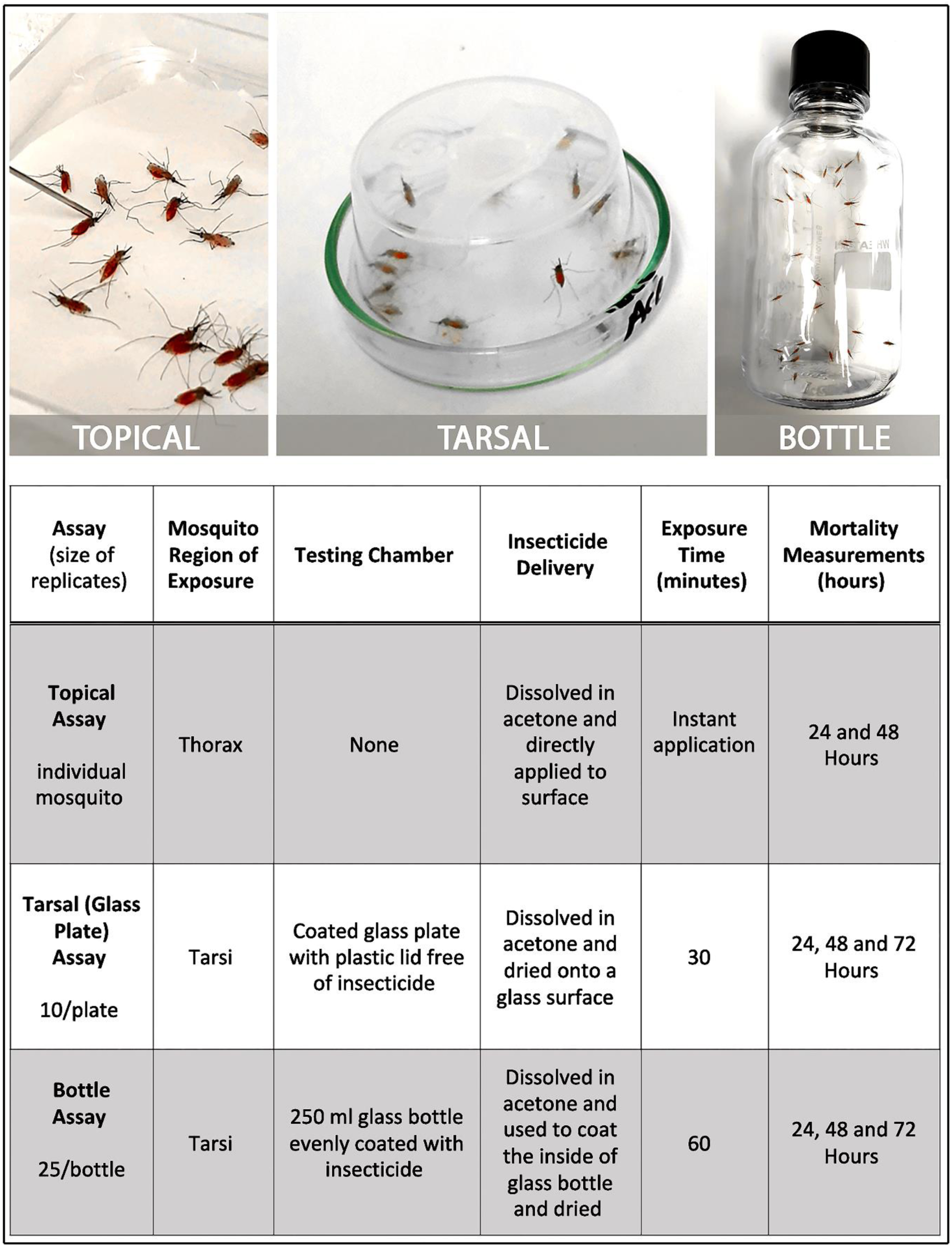
Comparison of the three assays used to assess the efficacy of nitisinone as a cuticular contact insecticide. Differences between the topical, tarsal and bottle assays, application method, insecticide delivery and time of exposure.

### Measuring the intrinsic activity of nitisinone through topical application at different stages of blood feeding

The intrinsic activity of nitisinone was measured by topically applying a nitisinone solution to the dorsal thorax of blood-fed mosquitoes (Kisumu) (Figure 6). Solutions of nitisinone were produced by dissolving nitisinone in acetone to produce concentrations of 1%, 0.1%, 0.01%, 0.001% and 0.0001%. (w/v). These solutions equate to 6, 0.6, 0.06, 0.006 and 0.0006 nmol/insect respectively. The concentrations were based on previous work used to screen insecticides against the Kisumu strain ^38^. Permethrin (Merck Life Science, Gillingham, UK), at 1% in acetone, was used as the positive control and the carrier, acetone, was the negative.

In groups of 60, mosquitoes were cold anaesthetised on ice for 10 mins. Once immobilised, the mosquitoes were oriented, so their dorsal thorax was facing up. A 0.2 µl drop of solution was applied to the dorsal thorax using a blunt end Hamilton syringe (Merck Life Science, Gillingham, UK). After dosing, mosquitoes were transferred to a large fabric cage (Bugdorm-4M3030 Watkins and Doncaster, Leominster, UK) to recover. The process was repeated with another set of 60 mosquitoes/concentration. Mortality was measured at 24h and 48h.

### Determination of discriminating dose using the WHO Bottle Assay

To determine the discriminating dose of nitisinone, a modification of the WHO Bottle assay was adopted ^48^ (Figure 6). Selected concentrations were established through dose determination assays (data not shown) and were selected to be 0, 15, 18, 20, 21.5, 22.5, 23.5, 25 and 30 µg per bottle with a 30 µg per bottle permethrin as a control. Nitisinone was dissolved in acetone at these concentrations and then added to 250 ml Wheaton Bottles (Fisher Scientific, Loughborough, UK).

Bottles were coated following previous methods ^38^ and allowed to dry overnight before use. To conduct the assay, 25 blood-fed female Kisumu mosquitoes (fed within 1 hr) were added to coated bottles and exposed for one hour. Mortality was scored at 1 h and 24 h. Three replicates were performed per concentration and the assay was repeated twice to total three biological replicates.

### Measuring the efficacy of nitisinone against susceptible and insecticide-resistant *An. gambiae* s.l., *C. quinquefasciatus* and *Ae. aegypti*

Using the glass plate bioassay previously described ^38^, a total of three replicates (biological) of thirty mosquitoes (ten mosquitoes per plate) were exposed to nitisinone-coated glass Petri dishes (radius 2.5cm, area 19.635cm^2^, SLS, Nottingham, UK). This assay compared the potency of nitisinone against blood-fed Kisumu, Tiassalé 13 and VK7 2014.

### Statistical Analysis

Graphs were constructed using GraphPad Prism (Version 9.4.1) except for the discriminating dose graph and probit analysis, which were constructed using PoloSuite 2.0 (LeOra Software). A Two-Way Anova (GraphPad) was used to assess the significance between data sets. Mortality was corrected using the Abbott’s Formula; if the negative control mortality was between 5-20%, a correction was made ^49^. There was no correction if mortality was below 5%. Replicates were discarded if mortality exceeded 20%.

### Image Production

All images were captured using a Nikon VVA241K001 1 J5 Compact System Camera (20.8 Mp) either handheld or mounted onto a Stemi 305 Stereo Microscope 8x-40x (Leica). Images were cropped, de- speckled, noise corrected, and size adjusted using Adobe PhotoShop CS (24.7.0 release).

## Data Availability

The datasets generated and/or analysed during the current study are available in the Mendeley Data repository, https://data.mendeley.com/preview/hs85hr246p?a=e83239b2-662d-40bf-b0b8-b88184729e86 (to be published upon acceptance)

## Acknowledgements

The authors wish to thank Dr Mary-Jo Hoare (LITE) for technical advice in assay design. Thank you to the technicians at the Liverpool Insect Testing Establishment for their help in rearing the resistant strains, particularly Ms Patricia McIntosh for aspirating the mosquitoes and organising delivery.

## Author Contributions

Z.S.D., L.R.H., R.S.L. and A.A.S. designed all the experiments. Z.S.D., G.P., C.R., F.B. and L.R.H. conducted experiments. Z.S.D. and L.R.H wrote the manuscript. All authors contributed to data interpretation, reviewed and edited the final version of the manuscript.

## Supplementary Data

**Figure S1.**
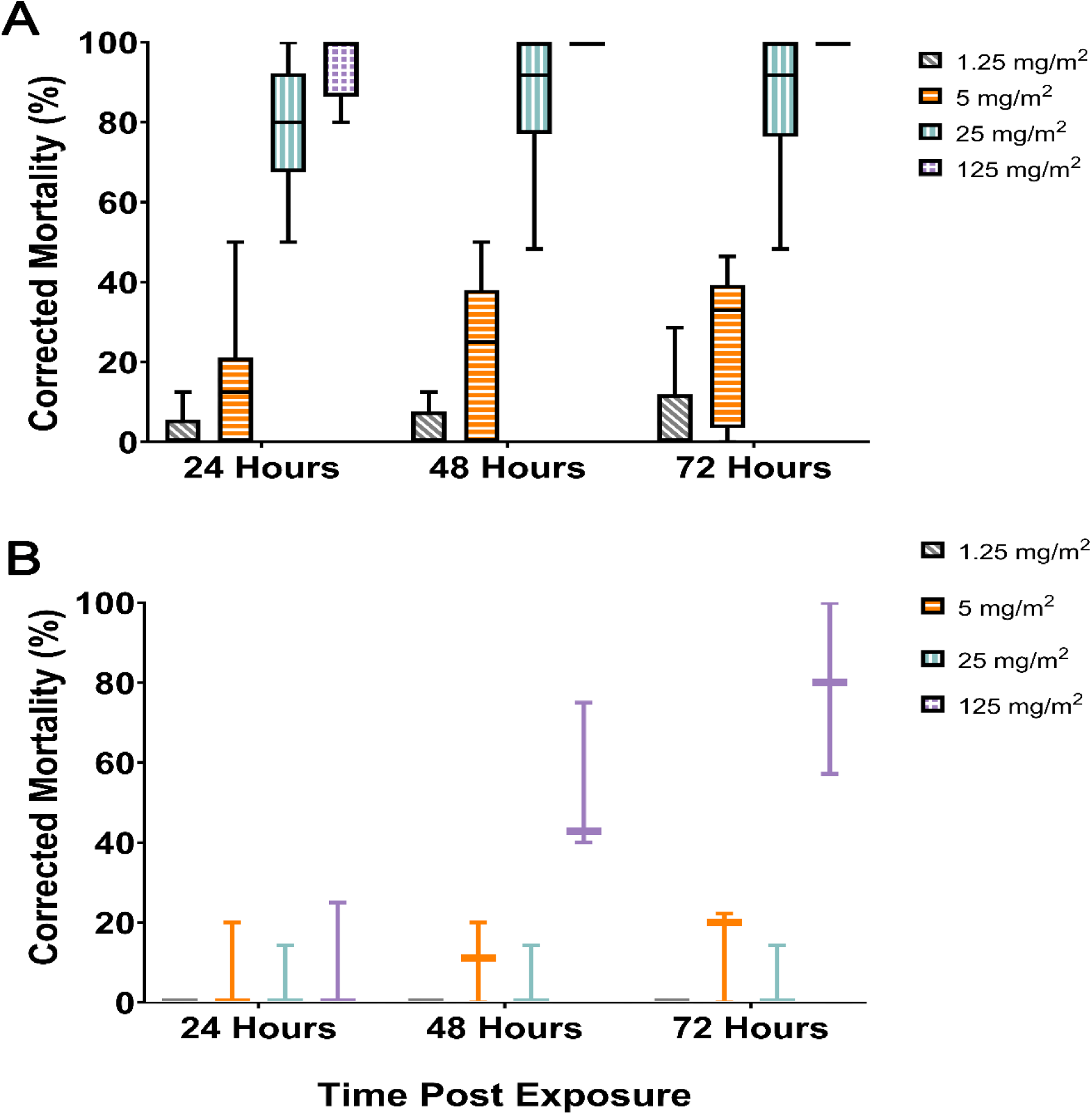
*Anopheles gambiae* Kisumu (A) and *Aedes aegypti* New Orleans (B) susceptibility profiles to tarsal contact with nitisinone prior to receiving a bloodmeal. A) Nitisinone concentrations tested: 1.25 mg/m^2^ (grey), 5 mg/m^2^ (orange), 25 mg/m^2^ (turquoise) and 125 mg/m^2^ (purple). Three biological replicates represent n = 90 per dose. B) 1.25 mg/m^2^ (grey), 5 mg/m^2^ (orange), 25 mg/m^2^ (turquoise) and 125 mg/m^2^ (purple). One biological replicate represent n = 30 per dose.

**Figure S2.**
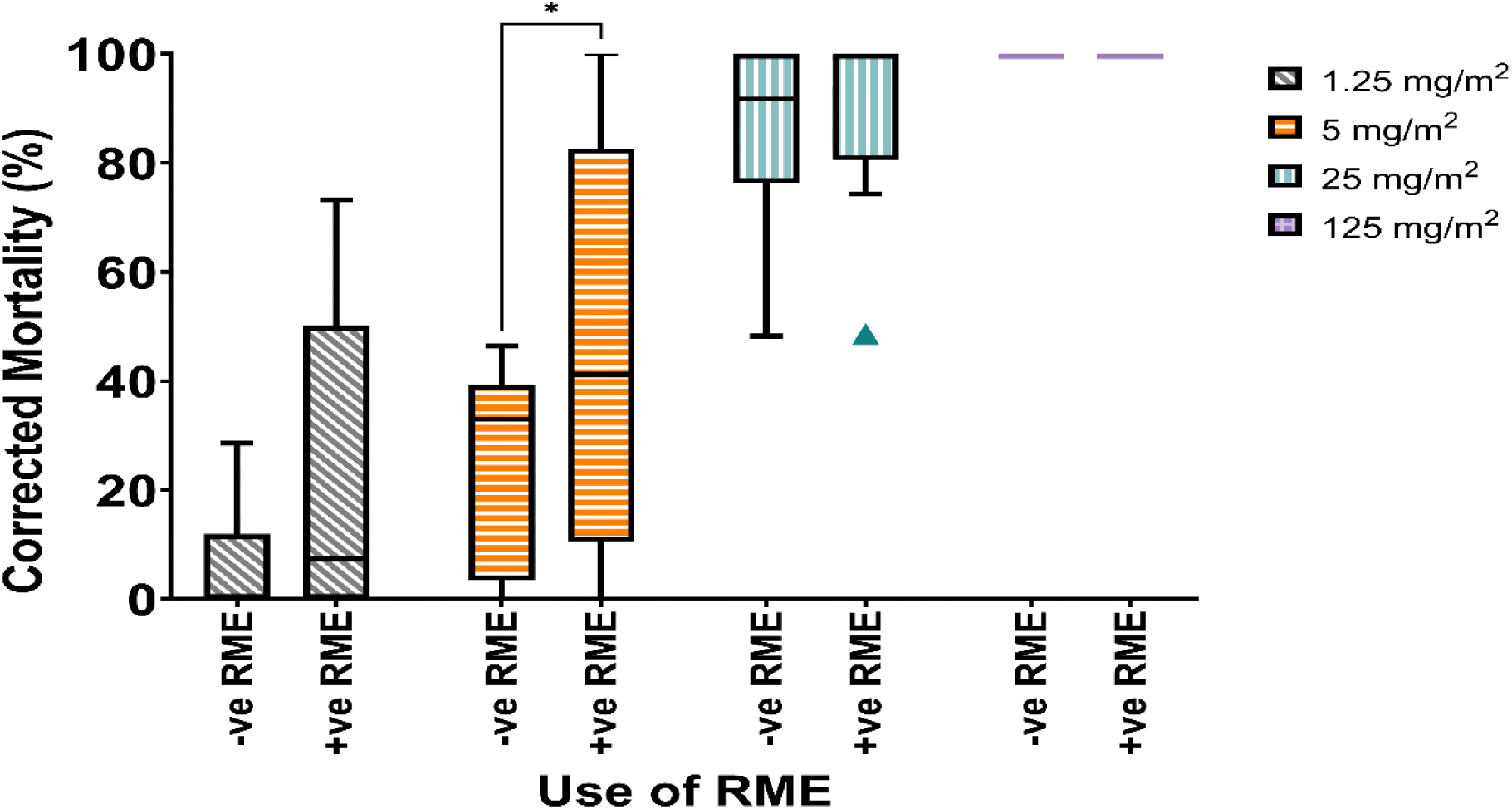
Anopheles gambiae Kisumu profile to tarsal contact with nitisinone prior to receiving a bloodmeal in the presence and absence of RME. Nitisinone concentrations tested: 1.25 mg/m2 (grey), 5 mg/m2 (orange), 25 mg/m2 (turquoise) and 125 mg/m2 (purple). RME was added at a concentration of 0.392 mg/ml. Three biological replicates represent n = 90 per dose.

**Table S1.** Insecticide resistance status of each mosquito strain used in this study. Status was determined through WHO Tube Assay and correct as of 2024. Insecticide classes are Pyrethroids (PY), organochlorides (OC), carbamates (CB) and organophosphates (OP).

| Species | Strain | Resistance Status |
| --- | --- | --- |
| *Anopheles gambiae* | Kisumu | Permethrin (PY): susceptible |
|  |  | Deltamethrin (PY): susceptible |
|  |  | Alpha-cypermethrin (PY): susceptible |
|  |  | DDT (OC): low levels of resistance detected |
|  |  | Dieldrin (OC): susceptible |
|  |  | Propoxur (CB): susceptible |
|  |  | Fenitrothion (OP): susceptible |
|  | Tiassalé 13 | Permethrin (PY): resistant |
|  |  | Deltamethrin (PY): resistant |
|  |  | Alpha-cypermethrin (PY): resistant |
|  |  | DDT (OC): resistant |
|  |  | Dieldrin (OC): resistant |
|  |  | Propoxur (CB): possible resistance |
|  |  | Fenitrothion (OP): susceptible |
|  | VK7 2014 | Permethrin (PY): resistant |
|  |  | Deltamethrin (PY): resistant |
|  |  | Alpha-cypermethrin (PY): resistant |
|  |  | DDT (OC): resistant |
|  |  | Dieldrin (OC): susceptible |
|  |  | Propoxur (CB): susceptible |
|  |  | Fenitrothion (OP): susceptible |
| *Aedes aegypti* | New Orleans | Permethrin (PY): susceptible |
|  |  | Deltamethrin (PY): susceptible |
|  |  | Alpha-cypermethrin (PY): susceptible |
|  |  | DDT (OC): susceptible |
|  |  | Dieldrin (OC): susceptible |
|  |  | Propoxur (CB): susceptible |
|  |  | Fenitrothion (OP): susceptible |
| *Culex quinquefasciatus* | Muheza | Permethrin (PY): resistant |
|  |  | Deltamethrin (PY): resistant |
|  |  | Alpha-cypermethrin (PY): resistant |
|  |  | DDT (OC): resistant |
|  |  | Dieldrin (OC): susceptible |
|  |  | Propoxur (CB): possible resistance |
|  |  | Fenitrothion (OP): susceptible |

**Table S2.** HPPD inhibitors used within this study. Compounds are given with their manufacturer and usage. Chemical and Toxicological information shown.

| Compound | Manufacturer | Use as Herbicide or Medicine | Solubility - In water at 20 °C (mg/L:) | Degradation Point | Mammals - Acute oral LD<sub>50</sub> (mg/kg) | Soil degradation (days) (aerobic) DT<sub>50</sub> (typical) |
| --- | --- | --- | --- | --- | --- | --- |
| Nitisinone | Swedish Orphan Biovitrum Ltd | Hereditary tyrosinaemia type I Alkaptonuria | 8.11 | n/a | n/a | n/a |
| Mesotrione | Greencrop Syngenta Chemsources | Grass and broad-leaved weeds | 1500 | 166 | >5000 | 19.6 |
| Sulcotrione | Syngenta | Grass and broad-leaved weeds | 165 | 170 | >5000 | 25 |
| Tembotrione | Bayer CropScience | Broad-leaved and grassy weeds | 71000 | 150 | >2500 | 14.5 |

